# Mechanisms of Cell Size Regulation in Slow-Growing *Escherichia coli* Cells: Discriminating Models Beyond the Adder

**DOI:** 10.1101/2023.09.11.557238

**Authors:** César Nieto, César Vargas-García, Juan Manuel Pedraza, Abhyudai Singh

**Affiliations:** Department of Physics, Universidad de los Andes, Bogotá, Colombia; Department of Electrical and Computing Engineering, University of Delaware. Newark, DE 19716, USA; Department of Systems Biology, Harvard Medical School, Boston, MA USA 02115; AGROSAVIA Corporación Colombiana de Investigación Agropecuaria. Mosquera. Colombia

## Abstract

Under ideal conditions, *Escherichia coli* cells divide after adding a fixed cell size, a strategy known as the *adder*. This concept applies to various microbes and is often explained as the division that occurs after a certain number of stages, associated with the accumulation of precursor proteins at a rate proportional to cell size. However, under poor media conditions, *E. coli* cells exhibit a different size regulation. They are smaller and follow a *sizer-like* division strategy where the added size is inversely proportional to the size at birth. We explore three potential causes for this deviation: precursor protein degradation, nonlinear accumulation rate, and a threshold size termed the *commitment size*. These models fit mean trends but predict different distributions given the birth size. To validate these models, we used the Akaike information criterion and compared them to open datasets of slow-growing *E. coli* cells in different media. the degradation model could explain the division strategy for media where cells are larger, while the commitment size model could account for smaller cells. The power-law model, finally, better fits the data at intermediate regimes.

## Introduction

To maintain tight distributions on size, bacteria must control the division time based on their size [1, 2]. Recent measurements indicate that bacteria, such as *Escherichia coli* and *Bacillus subtilis*, regularly divide using the *adder* division strategy [3, 4], where the size (Δ) added during a division cycle (birth to division) is not correlated with the size at birth (*s*_*b*_) [5, 6]. The molecular mechanisms behind *adder* division in bacteria are complex including the coordination of several processes, such as septal ring formation, DNA replication, and cell wall synthesis [4, 7–11]. A recent hypothesis suggests that a single factor, the accumulation of the FtsZ protein, is the main contributor to determining the timing of division under optimal growth conditions [4]. FtsZ forms a ring at the future division site and recruits other proteins to build a division apparatus [12]. Mathematical models describe the accumulation of FtsZ as a stochastic counting process with rates that depend on the size of the cell [13–15] opening new frontiers on the modelling of cell dynamics [16,17]. However, the exact dynamics of the FtsZ accumulation and the conditions under which it is the main contributor to the division remain unclear.

The *adder* mechanism may be broken under slow growth conditions [4, 18]. Then, Δ is negatively correlated with *s*_*b*_. This deviation from the *adder* is known as *sizer-like* strategy [4, 9, 18, 19] because it is an intermediate strategy between the *adder* and the *sizer*. In this last strategy, cells divide once they reach, on average, a specified size. This transition from *adder* to *sizer-like* by changing growth conditions could reveal more information on the division process and its regulation by different factors.

Here, we consider three competing models to explain these negative correlations: a) degradation of FtsZ [4, 20], which assumes that these molecules have a life span shorter than the doubling time; b) Non-linear size dependence of the FtsZ accumulation rate [18] and c) Additional size control mechanism [9, 21], which posits that FtsZ accumulation, and thus division, only starts when the cell reaches a minimal size. This commitment between cell cycle and a certain size is common in the analysis of the cell cycle for different microorganisms [22].

We observe that the three proposed models can fit the profile of Δ versus *s*_*b*_ [4, 18]. However, it is difficult to experimentally test which mechanism better determines the origin of the *sizer-like* in each particular condition, as it requires methods based on molecular biology that are not always easily accessible [4]. This research will present a data-based method in which, by comparing the predictions statistically with the data, it is possible to estimate which model explains the data with higher probability.

The paper is organized as follows: we first introduce the three models of bacterial division as special cases of a general stochastic counting process. We obtain the distributions of size at division by numerically solving the corresponding forward Kolmogorov equation for each model. We evaluate the models against the existing data sets [4, 18], using not only their mean trends but also their full distributions. We apply the Akaike Information Criterion (AIC), a likelihood-based method, to measure the fit of the models penalizing their complexity. We find that none of the models can explain the *sizer-like* strategy consistently, but some models perform better than others depending on how negative is the correlation Δ vs *s*_*b*_. Finally, we discuss the implications of our findings and suggest further experiments to investigate more aspects of cell division.

### Modelling division in rod-shaped bacteria

We consider the division process as the completion of a certain number of stages. During cell growth, as Figure 1 shows, division occurs exactly when a fixed number of stages *M* are crossed [14]. From a biological perspective, a possible interpretation of the division stages is related to the accumulation of a precursor protein, usually the FtsZ mentioned above [4, 23]. However, other molecules might also be the main contributor to bacterial divsion [24–27]. So, we use *division stages* instead of *number of precursor proteins* to explicitly keep our approach as general as possible. Note that *division stages* does not refer to checkpoints but rather to any mechanism that requires a certain number of sub-units to cause cell division.

**Figure 1.**
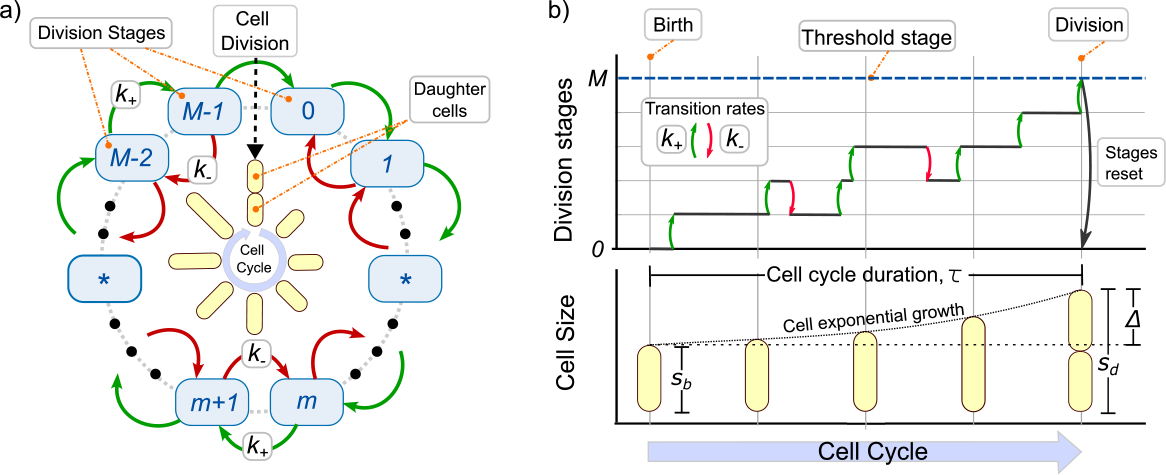
A general multistep model for triggering cell division. **(a)** Diagram of the cell cycle explaining how division occurs at crossing *M* stages. **(b)** While bacteria grow exponentially (lower panel), the division stages accumulate at rate *k*_+_ and might revert at rate *k−* (upper panel). Once a number of steps *M* is reached, the cell divides, the steps are reset to zero, and the size is halved. The main variables of the bacterial division cycle are also shown: size at birth *s*_*b*_, size at division *s*_*d*_, and added size Δ = *s*_*d*_ − *s*_*b*_ .

Figure 1 explains that a cell cycle is defined as the set of processes occurring during two consecutive divisions. During the cell cycle, the cell size *s* grows exponentially over time *t* with growth rate *μ*. This means that the cell size follows:

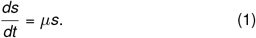

In the *adder* strategy, transition between stages occurs at a rate proportional to the current cell size [14, 18]. To explain the *sizer-like* strategy, we will generalize the accumulation rates. The rate of stage increase *k*_+_ is considered nonhomogeneous and, depending on the model, stages can revert (by protein degradation) with a rate *k*^*−*^. At cell birth, that is, at the beginning of the cell cycle, the cell starts from stage *m* = 0 and size *s*_*b*_ (which can differ from cell to cell). While cells grow exponentially, stages accumulate. When reaching the *m* = *M* stages, the cell divides. Exactly before division, the cell has a size *s*_*d*_ such that the added size Δ is defined as the difference Δ = *s*_*d*_−*s*_*b*_. Finally, during cell splitting, cell size is halved and the stages are reset to *m* = 0. To describe stage accumulation in the cell cycle, let *P*_*m*_(*t*) be the probability that *m* ≤ *M* stages will be completed at time *t* with *t* = 0 being the beginning of the cycle. Given the rates *k*_+_ and *k*^*−*^, the dynamics of these probabilities are described by the master equation:

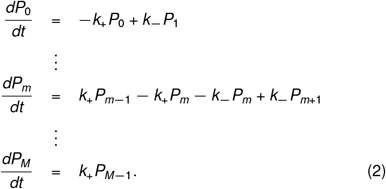

*P*_*M*_ is the probability of reaching the target step *M* or, equivalently, the probability of the division event to occur. Since after division the cell stars at stage *m* = 0, the initial condition (*t* = 0) is considered as *P*_0_ = *δ*_*m*,0_ with *δ*_*i,j*_ being the Kronecker delta.

As shown in Figure 2a, we are interested in the estimate of the time to division *τ*. *P*_*M*_ is related to *τ* as:

**Figure 2.**
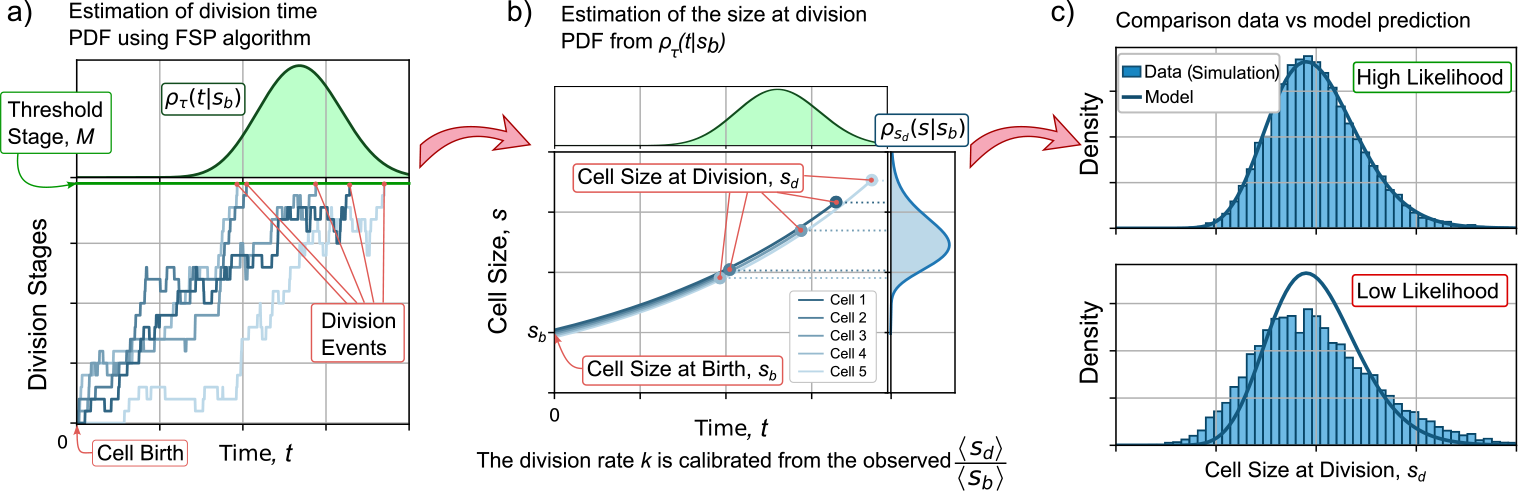
Process to predict the distribution of size at division and its comparison with data. **(a)** The distribution of division times *τ* given the size at birth *s*_*b*_ is estimated by solving numerically the master equation (2). **(b)** The size distribution at division *s*_*d*_ is obtained from the division times distribution and considering the exponential growth using (7). **(c)** The comparison with the data is made using methods based on likelihood. The distribution with higher likelihood fits the data better than a distribution with lower likelihood.

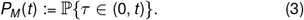

This is, *P*_*M*_ (*t*) is also the probability that *τ* occurs in the interval (0, *t*). Hence, the probability density function (also known as the distribution) of the division time *ρ*_*τ*_ is related to *P*_*M*_ following:

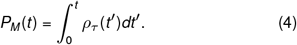

After estimating *P*_*M*_ by solving (2), *ρ*_*τ*_ (*t*) can be calculated as follows [28]:

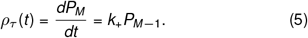

As shown in Figure 2b, the pdf of the cell size at division *s*_*d*_ can be estimated from *ρ*_*τ*_ considering that the cells grow exponentially. This is, the cell size *s* is related to the size at birth *s*_*b*_ through the time from birth *t* :

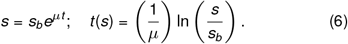

A transformation of variables allows us to obtain the pdf of sizes at division *ρ*_*s d*_ (*s*) as:

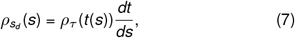

where 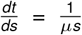 when exponential growth (1) is assumed. Observe how, since the cell size depends on *s*_*b*_, these distributions also depend on *s*_*b*_.

As explained in Figure 2b, a comparison between experiment and theory requires the calibration of the model parameters. This is done by estimating the moments of *s*_*d*_ using the estimated 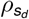 in (7). If ⟨ · ⟩ defines the averaging operator, the *α*-moment of the distribution of *s*_*d*_, written as 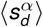is defined as follows,

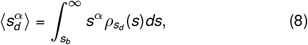

where the mean size at division ⟨*s*_*d*_⟩, the moment with *α* = 1, can be used to calculate the mean added size per division cycle ⟨Δ⟩ = ⟨*s*_*d*_⟩−*s*_*b*_ as a function of the size at birth *s*_*b*_. The particular parameters of the model (as will be explained later) are adjusted such as the predicted ⟨*s*_*d*_⟩ is constrained to the observed average *s*_*d*_ assuming that the mean size at birth⟨*s*_*b*_⟩ = 1.

After imposing this constraint, the best model parameters are adjusted to the data by maximizing the likelihood function. As shown in Figure 2c, the likelihood function measures the precision of the pdf with the histogram associated with the data. With a higher likelihood, the predicted distribution fits the experiment better.

The moments of *s*_*d*_ can also be used to quantify the noise in added size 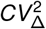 the ratio between var(Δ) (the variance of Δ) and ⟨Δ⟩^2^, which is a measure of stochastic variability Δ. We^Δ^can obtain 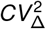 as a function of *s*_*b*_ from 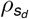 and its moments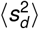and ⟨*s*_*d*_ ⟩ using the formula [18]:

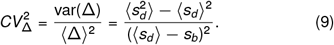

Observe how since 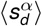 depends on *s*_*b*_, different models may predict different trends on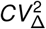. For us, while the trend ⟨Δ ⟩ vs *s*_*b*_ is known as the division strategy, 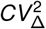 vs *s*_*b*_ is the noise signature of the model. Next, we will explain these models as particular cases of *k*_+_ and *k*^*−*^.

### The *Adder* strategy

The implementation of the *adder*, where ⟨Δ⟩ is independent on *s*_*b*_ corresponds to the particular case of (2) where *k*_+_ and *k*^*−*^ are given by [18]:

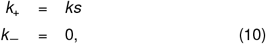

with *k* a constant and *s* = *s*_*b*_*e*^*μt*^ is the cell size. Assuming exponentially growing cells and division process defined by both (2) and (10), the mean added cell size ⟨Δ⟩ is given by [18]:

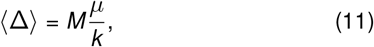

which is, as expected, independent of *s*_*b*_. The noise in Δ, defined in (9) follows:

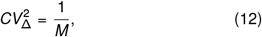

which is also uncorrelated with *s*_*b*_ as observations suggest [18].

### *Sizer-like* by molecule degradation

As [4] suggests, *sizer-like* division strategy can be obtained by including active degradation of the division triggering molecules. In our framework, degradation is equivalent to a step backward in the accumulation of *M* stages. In this case, the rates *k*_+_ and *k*^*−*^ are given by:

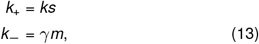

where γ is the rate at which each molecule is degraded

A negative slope is obtained in Δ vs *s*_*b*_ as shown in Figure 3a (top). The higher *γ*, the more pronounced the slope. This model reduces to the *adder* when *γ* ≪*μ*. The noise signature of this model is presented in Figure 3a (bottom) where, as the main property, we can see that an increase *γ*, for fixed division steps *M*, is expected a higher average 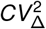 .

**Figure 3.**
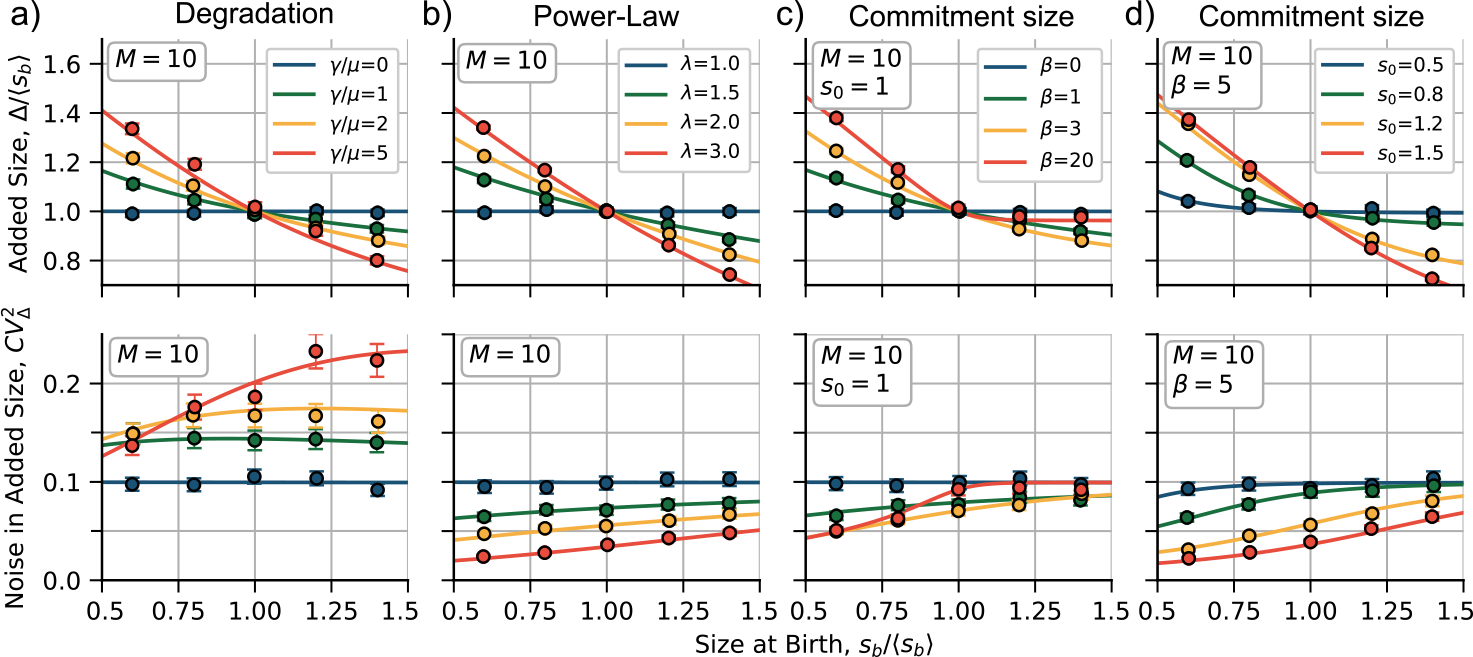
Trends on mean added size before division ⟨Δ⟩ **(top) and its stochastic fluctuations (noise)** 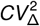**(bottom) as functions of the size at birth** *s*_*b*_ . **(a)** Trends considering the model of degradation for different values of the degradation rate *γ* relative to the growth rate *μ* in (13). **(b)** Predictions considering a division rate proportional to a power of size for different exponents *λ* in (14). **(c)** and **(d)** are the predictions consideringa commitment size with different values of the Hill function exponent *β* and the commitment size *s*_0_, respectively, in (15). The trend lines correspond to the numerical solutions of (2). Large dots are obtained from simulations. Error bars represent the 95%-confidence interval over 10000 simulated cycles. The constant *k* is set in each case such as ⟨Δ⟩ = ⟨*s*_*b*_ ⟩ = 1. Other parameters are shown inset.

### Sizer-like via non-linear division rate

Following [18], we consider a scenario where a molecule that triggers division is produced at a rate that depends on a power *λ* of cell size *s*, this is:

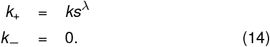

After substituting (14) in (2), plus the assumption of exponential growth, the division strategy exhibits a *sizer-like* behavior when *λ >* 1 (Figure 3b top). For the particular case of *λ* = 1, the model reduces to the *adder*. Hence, the higher *λ*, the higher the slope of the relationship between Δ and *s*_*b*_. The power-law on the production rate also affects the fluctuations of the added size 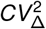: it lowers the average noise level for a given *M* and simultaneously increases the positive slope of 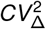 versus *s*_*b*_ (Figure 3b bottom).

### Sizer-like via a commitment size

Recent evidence suggests that cells may aim for a minimal size before starting division programs [9]. We will denote this minimal or commitment size by *s*_0_. We propose that it can be incorporated into our framework as

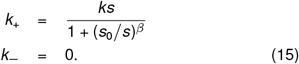

By replacing (15) with (2), we can see that the division strategy has a *sizer-like* behavior when *β >* 0 (Figure 3c top). The parameter *β* can control the slope of the curve Δ versus *s*_*b*_ for a fixed *s*_0_. It also influences the fluctuations of the added size 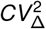 : It reduces the average noise level for a given *M*, but increases the positive slope of 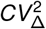 versus *s*_*b*_ (Figure 3c bottom). This model encompasses the *adder* strategy as a special case when *β* = 0.

The commitment size *s*_0_ relative to the mean size at birth⟨*s*_*b*_⟩ also affects the division strategy. For a fixed *β*, a low *s*_0_≪*s*_*b*_ mimics the *adder*, while a high *s*_0_≫*s*_*b*_ approximates the *sizer* (the strategy where the slope in Δ versus *s*_*b*_ is -1). This is because cells born with a size below *s*_0_ follow a perfect *sizer* strategy, while cells born with size above *s*_0_ follow the *adder*. For small *β*, the *adder* strategy is recovered. For intermediate *β*, the transition from *sizer* to *adder* is smoother as *s*_*b*_ increases.

### Comparison theory vs datasets

We have shown how to derive the pdf *ρ*(*s*_*d*_|*s*_*b*_) from different models. Now, we want to estimate how accurate these distributions are relative to the data. We use the Akaike Information Criterion (AIC) [29] to reward the fit of the models to the data, while penalizing the number of free parameters. The adder model has one parameter (*M*), the degradation and power-law models have two (*γ, M* and *λ, M* respectively), and the commitment size model has three (*s*_0_, *β* and *M*). Taking into account all experiments, for each pair of data (*s*_*b*_, Δ) normalized by ⟨*s*_*b*_⟩ we numerically compute the size-at-division distribution 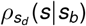 given *s*_*b*_ using (7). We find the parameters that maximize the likelihood function [30] based on the data. Then we calculate the AIC for each model. The model with the lowest AIC is the most probable one, and we denote its AIC value by *AIC*_*min*_. To compare the relative probability of each model with the most probable one, we use the concept of relative likelihood [31]. For a model *i* with an AIC value of *AIC*_*i*_, the relative likelihood *p* is given by:

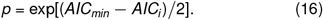

The AIC method is useful for estimating the accuracy of the models, but is not easy to visualize since the match of the distributions is very similar. To gain a better intuitive understanding of how each model behaves, we can use the method of statistical moments. As shown in Figure 3, this method involves plotting the division strategy (⟨Δ⟩versus *s*_*b*_) and the noise signature (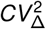versus ⟨*s*_*b*_⟩) and comparing them with the data. From theory, we can obtain the moments directly: given a *s*_*b*_, they are calculated from the distribution using (8). From the data set, we visualize the moments given *s*_*b*_ using quantile splitting. This method splits the data into a given number of quantiles and computes the statistics for each quantile separately. The points in Figure 3 (five quantiles) represent the data from simulations, while Figure 4a (also five quantiles) represents the experimental data. To study the division strategy, we plot ⟨*s*_*b*_⟩ _*q*_ and ⟨Δ⟩ _*q*_ for each quantile. The noise signature is obtained by plotting the variance 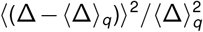 for each quantile. Error bars indicate a 95% confidence interval using bootstrapping methods.

**Figure 4.**
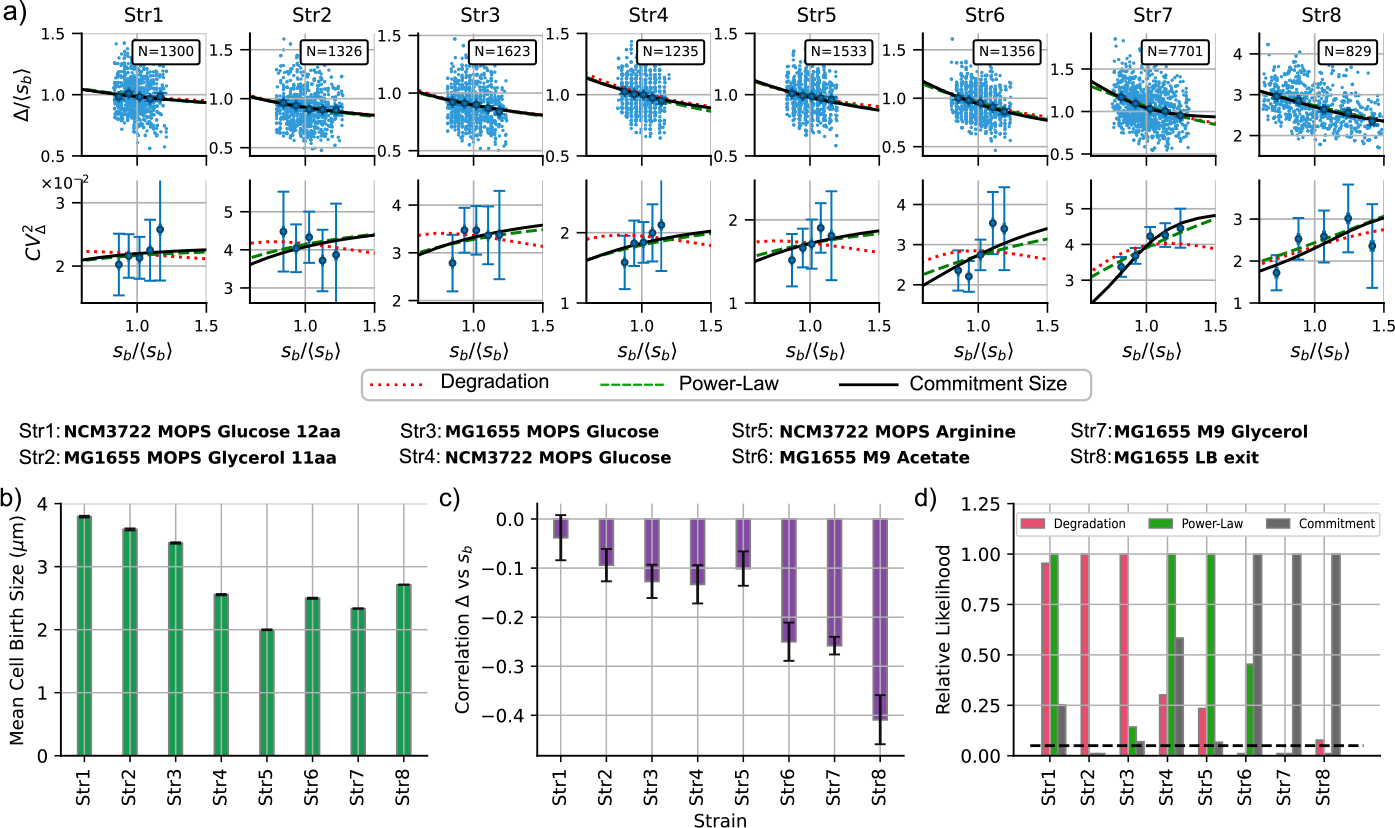
Discriminating between models across different experimental conditions. **(a)** Top: trends of the added size Δ vs the size at birth *s*_*b*_. Bottom: Noise signatures, as quantified by the stochastic fluctuations of added size 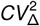 vs *s*_*b*_. The numerical prediction of the three models can be discriminated better: degradation model (red dotted line), power law (green dashed line), and commitment size (black solid line). *N* represents the number of studied cell cycles. Str represents the strain number. In the middle of the figure, more detailed labels are shown. **(b)** Comparison of the mean cell size at birth for the studied strains. **(c)** Comparison of the correlation function between Δ and *s*_*b*_ for strains. The more negative this correlation is, the closer to *sizer* the division strategy is. **(d)** Relative likelihood of each model with respect to the model with the best AIC score using (16). The black dashed line represents the relative likelihood of 0.05.

### Experiments

We analyzed three independent already published datasets of *E. coli* strains under different growth conditions [4, 18, 32]. Data were obtained from time-lapse microscopy images of single cells confined and fed in a *mother machine* micro-fluid device. References [4, 18] imaged slow-growing cells corresponding to steady growth conditions (strains 1 to 7), while [32] studied the first cell division of cells resuspended in rich growth medium (strain 8).

We measured the cell size using the cell length since the width is approximately constant. We normalize all lengths by the mean size at birth ⟨*s*_*b*_⟩. The theoretical division rate *k* was estimated, given the free parameters and setting the growth rate to *μ* = ln(2), from the observed ⟨Δ⟩ with⟨*s*_*b*_⟩ = 1. The other properties of cell division such as the slope in Δ vs. *s*_*b*_ or 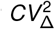 vs. *s*_*b*_ emerge naturally given the model parameters. The experimental data and data analysis scripts are available at [33].

## Results

Figure 4a shows the comparison between the data and the theory. First, we study the case where the division strategy is *adder*, that is, when Δ is independent of *s*_*b*_. This occurs for strain1 (NCM3722 in MOPS with Glucose) since the 95% confidence interval for the correlation includes zero (Figure 4c). For these conditions, the three models reach a similar likelihood, but the commitment size model is punished for its complexity. Both degradation a power-law reach similar AIC score and is not possible to discard one of those models with enough confidence.

For the other strains, the division is *sizer-like* with statistical significance (Figure 4c). In Figure 4d, we observe that the degradation model has a minimum AIC (and therefore highest relative likelihood) for cells with a larger mean size at birth (Strains 2 and 3, MG1655 in MOPS with glucerol 11aa and MOPS with Glucose). For smaller strains, power-law and commitment show lower AICs. While powerlaw has the lowest AIC forstrains 4 and 5 (NCM3722 in MOPS glucose and MOPS arginine), the commitment size model performs best for strains with the lowest correlation Δ vs *s*_*b*_ (strains 6, 7 and 8: MG1655 in M9 with Acetate and M9 with Glycerol and first division in LB).

To better visualize the performance of the models, we can compare the statistical moments with the binned moments of the data. In Figure 4a, we observe that all of our proposed models capture the mean trend in Δ vs. *s*_*b*_. However, they differ in the noise signature 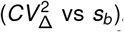. The degradation model predicts a low correlation between [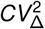 and *s*_*b*_, while the power-law model predicts an increasing function. The commitment size model behaves like the power law when the slope of Δ vs. *s*_*b*_ is small, but predicts a higher slope of 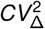 vs *s*_*b*_ when the slope of Δ vs. *s*_*b*_ is large.

## Discussion

The transition from the *adder* to the *sizer-like* division strategies suggests the presence of a dual mechanism governing cell division: a primary mechanism that leads to the *adder* being the most important under optimal fast-growth conditions and larger cells, and a secondary mechanism that gives rise to the *sizer-like* is more visible in suboptimal slow-growth conditions and smaller cells. It is plausible that this secondary mechanism has often been overlooked in laboratory settings, where cells are commonly cultivated in nutrient-rich environments. However, this mechanism is of substantial importance, as it could elucidate cell survival and adaptation in real-world scenarios characterized by less-than-optimal or slower growth conditions.

Si *et al*. [4] found experimental evidence that the degradation of FtsZ could underlie the *sizer-like* strategy, supported by the restoration of *adder* division in *E. coli* upon inhibiting FtsZ degradation through *clpX* knockdown. Our investigation highlights the potency of the degradation model in explaining the *sizer-like* strategy, particularly when the relationship between added size and birth size is not excessively steep. However, for steeper slopes, alternative mechanisms align better with the data. We believe that the actual mechanism explaining the *sizer-like* is a composition of degradation and commitment size, since they are not exclusive.

The commitment size model posits a minimum size prerequisite for division initiation, which yields the *sizer-like* behavior. In environments characterized by slow growth or limited nutrents, cells are smaller, often falling below this commitment size, needing to grow until reaching this threshold. Then the division machinery starts to build up. In contrast, under nutrient-rich conditions, cells at birth exceed the commitment size, prompting division immediately. With division starting just after birth, we expect large cells to follow the *adder* strategy. Empirical support for this concept emerges from experimental observations (strain 8 in Figure 1). In this strain, bacteria grown overnight are approximately one-third the size of cells under steady growth conditions. The first division post-resuspension shows a strong *sizer-like* strategy, while subsequent divisions, where cells are larger, adopt the conventional *adder* division [32]. The commitment size model offers the best fit for these data using AIC metrics.

The power-law rate model operates on the premise that the division rate has a stronger dependence on the size that the *adder* model. It is important to note that this model is heuristic, using nonlinear size dependency as an effective parameter to capture the intricacies of division. Thus, the power-law rate may approximate the Hill function rate of the commitment size model within a specific range. In contrast, the commitment size model, with more free parameters, can exhibit greater adaptability compared to the power-law model.

The notion of commitment size might relate to the initiation of chromosome replication at a fixed size per origin [9, 34, 35]. Although our model does not explicitly integrate chromosome replication, potential links between these variables remain open. Recent studies hint at coordination between division and replication in bacteria, resembling observations in more complex organisms [27, 36–38]. However, there is evidence that replication and division are essentially weakly-coupled processes since division is related to the accumulation of FtsZ and replication is related with the protein DnaA [4].

The underlying biological mechanism enabling the cell to measure the commitment size remains a subject of active debate. One perspective suggests tight synchronization between division and DNA replication, which triggers division when reaching a target size for replication initiation [9, 39, 40]. Another angle posits the existence of molecules acting as size proxies within cells, regulating division initiation [41,42]. Further experiments, as proposed in [32, 43], could provide additional information on the division strategy. Dynamic environments, where the division strategy changes dynamically, could shed light on these mechanisms. Other ways to achieve slow-growth conditions can also help us understand the validity of the models. These conditions can include decreasing growth temperature and growing in the presence of a mild concentration of antibiotics. Furthermore, exploing the division strategy with the *clpX* knockdown strain under different growth conditions, coupled with the use of AIC or other likelihood-based methods for data analysis, has the potential to provide a more comprehensive insight into these phenomena.

## Author Contributions

CN and CVG developed the model, CN analyzed the data and designed the plots. CVG, JMP and AS guided the research. All the authors contributed on the writing of the article.

## Acknowledgments

CN was supported by Colombian Science Ministry *Convocatoria 647 para doctorados nacionales*. AS acknowledges support from NIH-NIGMS via grant R35GM148351.

## References

[1] C. A. Vargas-Garcia, M. Soltani, and A. Singh, “Conditions for cell size homeostasis: a stochastic hybrid system approach,” IEEE Life Sciences Letters, vol. 2, no. 4, pp. 47–50, 2016.

[2] S. Taheri-Araghi, S. Bradde, J. T. Sauls, N. S. Hill, P. A. Levin, J. Paulsson, M. Vergassola, and S. Jun, “Cellsize control and homeostasis in bacteria,” Current Biology, vol. 25, no. 3, pp. 385–391, 2015.

[3] S. Jun and S. Taheri-Araghi, “Cell-size maintenance: universal strategy revealed,” Trends in microbiology, vol. 23, no. 1, pp. 4–6, 2015.

[4] F. Si, G. Le Treut, J. T. Sauls, S. Vadia, P. A. Levin, and S. Jun, “Mechanistic origin of cell-size control and homeostasis in bacteria,” Current Biology, vol. 29, no. 11, pp. 1760–1770, 2019.

[5] S. Jun, F. Si, R. Pugatch, and M. Scott, “Fundamental principles in bacterial physiology—history, recent progress, and the future with focus on cell size control: a review,” Reports on Progress in Physics, vol. 81, no. 5, p. 056601, 2018.

[6] M. Campos, I. V. Surovtsev, S. Kato, A. Paintdakhi, B. Beltran, S. E. Ebmeier, and C. Jacobs-Wagner, “A constant size extension drives bacterial cell size homeostasis,” Cell, vol. 159, no. 6, pp. 1433–1446, 2014.

[7] P. Kar, S. Tiruvadi-Krishnan, J. Männik, J. Männik, and A. Amir, “Using conditional independence tests to elucidate causal links in cell cycle regulation in escherichia coli,” Proceedings of the National Academy of Sciences, vol. 120, no. 11, p. e2214796120, 2023.

[8] J. Männik, B. E. Walker, and J. Männik, “Cell cycledependent regulation of ftsz in escherichia coli in slow growth conditions,” Molecular microbiology, vol. 110, no. 6, pp. 1030–1044, 2018.

[9] M. Wallden, D. Fange, E. G. Lundius, Ö. Baltekin, and J. Elf, “The synchronization of replication and division cycles in individual e. coli cells,” Cell, vol. 166, no. 3, pp. 729–739, 2016.

[10] M. Priestman, P. Thomas, B. D. Robertson, and V. Shahrezaei, “Mycobacteria modify their cell size control under sub-optimal carbon sources,” Frontiers in cell and developmental biology, vol. 5, p. 64, 2017.

[11] H. E. Opalko, K. E. Miller, H.-S. Kim, C. A. Vargas-Garcia, A. Singh, M.-C. Keogh, and J. B. Moseley, “Arf6 anchors cdr2 nodes at the cell cortex to control cell size at division,” Journal of Cell Biology, vol. 221, no. 2, p. e202109152, 2021.

[12] H. P. Erickson, D. E. Anderson, and M. Osawa, “Ftsz in bacterial cytokinesis: cytoskeleton and force generator all in one,” Microbiology and molecular biology reviews, vol. 74, no. 4, pp. 504–528, 2010.

[13] C. Jia, A. Singh, and R. Grima, “Cell size distribution of lineage data: analytic results and parameter inference,” Iscience, vol. 24, no. 3, p. 102220, 2021.

[14] K. R. Ghusinga, C. A. Vargas-Garcia, and A. Singh, “A mechanistic stochastic framework for regulating bacterial cell division,” Scientific reports, vol. 6, p. 30229, 2016.

[15] M. Basan, M. Zhu, X. Dai, M. Warren, D. Sévin, Y.-P. Wang, and T. Hwa, “Inflating bacterial cells by increased protein synthesis,” Molecular systems biology, vol. 11, no. 10, p. 836, 2015.

[16] C. Nieto, S. C. Blanco, C. Vargas-García, A. Singh, and P. J. Manuel, “Pyecolib: a python library for simulating stochastic cell size dynamics,” Physical Biology, vol. 20, no. 4, p. 045006, 2023.

[17] D. Serbanescu, N. Ojkic, and S. Banerjee, “Cellular resource allocation strategies for cell size and shape control in bacteria,” The FEBS Journal, vol. 289, no. 24, pp. 7891–7906, 2022.

[18] C. Nieto, J. Arias-Castro, C. Sánchez, C. Vargas-García, and J. M. Pedraza, “Unification of cell division control strategies through continuous rate models,” Physical Review E, vol. 101, no. 2, p. 022401, 2020.

[19] J. T. Sauls, D. Li, and S. Jun, “Adder and a coarse-grained approach to cell size homeostasis in bacteria,” Current opinion in cell biology, vol. 38, pp. 38–44, 2016.

[20] K. Sekar, R. Rusconi, J. T. Sauls, T. Fuhrer, E. Noor, J. Nguyen, V. I. Fernandez, M. F. Buffing, M. Berney, S. Jun, et al., “Synthesis and degradation of ftsz quantitatively predict the first cell division in starved bacteria,” Molecular systems biology, vol. 14, no. 11, p. e8623, 2018.

[21] H. Shi, Y. Hu, P. D. Odermatt, C. G. Gonzalez, L. Zhang, J. E. Elias, F. Chang, and K. C. Huang, “Precise regulation of the relative rates of surface area and volume synthesis in bacterial cells growing in dynamic environments,” Nature communications, vol. 12, no. 1, p. 1975, 2021.

[22] A. Litsios, D. H. Huberts, H. M. Terpstra, P. Guerra, A. Schmidt, K. Buczak, A. Papagiannakis, M. Rovetta, J. Hekelaar, G. Hubmann, et al., “Differential scaling between g1 protein production and cell size dynamics promotes commitment to the cell division cycle in budding yeast,” Nature cell biology, vol. 21, no. 11, pp. 1382–1392, 2019.

[23] L. K. Harris and J. A. Theriot, “Relative rates of surface and volume synthesis set bacterial cell size,” Cell, vol. 165, no. 6, pp. 1479–1492, 2016.

[24] Q. Zhang, Z. Zhang, and H. Shi, “Cell size is coordinated with cell cycle by regulating initiator protein dnaa in e. coli,” Biophysical Journal, vol. 119, no. 12, pp. 2537–2557, 2020.

[25] C. Speck and W. Messer, “Mechanism of origin unwinding: sequential binding of dnaa to double-and single-stranded dna,” The EMBO journal, vol. 20, no. 6, pp. 1469–1476, 2001.

[26] G. Schreiber, E. Z. Ron, and G. Glaser, “ppgpp-mediated regulation of dna replication and cell division in escherichia coli,” Current microbiology, vol. 30, no. 1, pp. 27–32, 1995.

[27] G. Micali, J. Grilli, M. Osella, and M. C. Lagomarsino, “Concurrent processes set e. coli cell division,” Science Advances, vol. 4, no. 11, p. eaau3324, 2018.

[28] K. R. Ghusinga, J. J. Dennehy, and A. Singh, “First-passage time approach to controlling noise in the timing of intracellular events,” Proceedings of the National Academy of Sciences, vol. 114, no. 4, pp. 693–698, 2017.

[29] Y. Sakamoto, M. Ishiguro, and G. Kitagawa, “Akaike information criterion statistics,” Dordrecht, The Netherlands: D. Reidel, vol. 81, no. 10.5555, p. 26853, 1986.

[30] T. A. Severini, Likelihood methods in statistics. Oxford University Press, 2000.

[31] K. P. Burnham and D. R. Anderson, “Multimodel inference: understanding aic and bic in model selection,” Sociological methods & research, vol. 33, no. 2, pp. 261–304, 2004.

[32] S. Bakshi, E. Leoncini, C. Baker, S. J. Cañas-Duarte, B. Okumus, and J. Paulsson, “Tracking bacterial lineages in complex and dynamic environments with applications for growth control and persistence,” Nature Microbiology, vol. 6, no. 6, pp. 783–791, 2021.

[33] C. Nieto, “Sizer-like division analysis. doi: 10.5281/zenodo.3951080,” Sept. 2023.

[34] P.-Y. Ho and A. Amir, “Simultaneous regulation of cell size and chromosome replication in bacteria,” Frontiers in microbiology, vol. 6, p. 662, 2015.

[35] J. Chen, H. Boyaci, and E. A. Campbell, “Diverse and unified mechanisms of transcription initiation in bacteria,” Nature Reviews Microbiology, vol. 19, no. 2, pp. 95–109, 2021.

[36] X.-M. Sun, A. Bowman, M. Priestman, F. Bertaux, A. Martinez-Segura, W. Tang, C. Whilding, D. Dormann, V. Shahrezaei, and S. Marguerat, “Size-dependent increase in rna polymerase ii initiation rates mediates gene expression scaling with cell size,” Current Biology, vol. 30, no. 7, pp. 1217–1230, 2020.

[37] N. E. Kleckner, K. Chatzi, M. A. White, J. K. Fisher, and M. Stouf, “Coordination of growth, chromosome replication/segregation, and cell division in e. coli,” Frontiers in microbiology, vol. 9, p. 1469, 2018.

[38] N. Ramkumar and B. Baum, “Coupling changes in cell shape to chromosome segregation,” Nature Reviews Molecular Cell Biology, vol. 17, no. 8, pp. 511–521, 2016.

[39] H. Zheng, Y. Bai, M. Jiang, T. A. Tokuyasu, X. Huang, F. Zhong, Y. Wu, X. Fu, N. Kleckner, T. Hwa, et al., “General quantitative relations linking cell growth and the cell cycle in escherichia coli,” Nature Microbiology, vol. 5, no. 8, pp. 995–1001, 2020.

[40] A. Knöppel, O. Broström, K. Gras, J. Elf, and D. Fange, “Regulatory elements coordinating initiation of chromosome replication to the escherichia coli cell cycle,” Proceedings of the National Academy of Sciences, vol. 120, no. 22, p. e2213795120, 2023.

[41] G. Le Treut, F. Si, D. Li, and S. Jun, “Quantitative examination of five stochastic cell-cycle and cell-size control models for escherichia coli and bacillus subtilis,” Frontiers in microbiology, p. 3278, 2021.

[42] M. Berger and P. R. t. Wolde, “Robust replication initiation from coupled homeostatic mechanisms,” Nature communications, vol. 13, no. 1, p. 6556, 2022.

[43] C. Nieto, C. Vargas-García, J. M. Pedraza, and A. Singh, “Modeling cell size control under dynamic environments,” IFAC-PapersOnLine, vol. 55, no. 40, pp. 133–138, 2022.

