## Supplementary material for "Mechanisms of Cell Size Regulation in Slow-Growing *Escherichia coli* Cells: Discriminating Models Beyond the Adder": DataAnalysis.ipynb

```
# %% [code]
import pandas as pd
import numpy as np
from matplotlib import pyplot as plt
import scipy
from scipy.stats import bayes_mvs as bayesest
import os #Lib for get the operative system sintaxis
from sklearn import linear_model#Machine learning package for a smart plot fit
from scipy import stats
from sklearn.metrics import mean_squared_error as msqerr
from tqdm import tqdm
from scipy import optimize
import random

# %% [code]

def bootstrap(arr):
    #Function for estimating the mean and its 95% confidence interval using boostraping
    mnar=np.empty(1000)
    for l in range(1000):#Selecting 1000 random datasets from the array 
        mn=np.mean(random.choices(arr,k=len(arr)))
        mnar[l]=mn
    mn=np.mean(mnar)
    up=np.quantile(mnar,0.975)
    down=np.quantile(mnar,0.025)
    return(mn,0.5*(-down+up))

def bootstrapcv2(arr):
    #Function for estimating the CV squared and its 95% confidence interval using boostraping methods
    mnar=np.empty(1000)
    for l in range(1000):
        xx=random.choices(arr,k=len(arr))
        mn=np.var(xx)/np.mean(xx)**2
        mnar[l]=mn
    mn=np.mean(mnar)
    up=np.quantile(mnar,0.975)
    down=np.quantile(mnar,0.025)
    return(mn,0.5*(-down+up))
def bootcorr(x,y):
    #Function for estimating the correlation function of two dataarrays 
    #squared and its 95% confidence interval using boostraping
    slopearr=[]
    for i in range(1000):
        xboot=[]
        yboot=[]
        for m in range(len(x)):
            i=np.random.randint(0,len(x))
            xboot.append(x[i])
            yboot.append(y[i])
        corr=np.corrcoef(xboot,yboot)[0,1]
        slopearr.append(corr)
    mn=np.mean(slopearr)
    up=np.quantile(slopearr,0.975)
    down=np.quantile(slopearr,0.025)
    return(mn,0.5*(-down+up))

# %% [code]
def setaxis(ax):
    for axis in ['bottom','left']:
        ax.spines[axis].set_linewidth(1)
    for axis in ['top','right']:
        ax.spines[axis].set_linewidth(0)
    ax.grid()
    ax.tick_params(axis='both', labelsize=8,length=5)

# %% [code]
def getcum(sb,steps,gammap,s0h,H,lamb,k):
    """
    This function returns the probabilty density function for the size at division and the range over 
    which it is defined
    :Param float size at birth relative to the mean size at birth (non-negative)
    :Param int Steps: Division steps to trigger division (Positive)
    :Params float gammap: Degradation rate (non-negative)
    :param float s0h: Comitment size (non-negative)
    :param float H: Hill function coefficient (non-negative)
    :param float lamb: exponent of power law for division rate (non-negative)
    :param float k: Division rate
    """ 
    
    kk=k
    mu=np.log(2)

    def du(u,sb,t):
        mu=np.log(2)
        v=np.zeros_like(u)
        s=sb*np.exp(mu*t)
        k=kk*kd(s=s,H=H,s0h=s0h)
        for l in range(len(u)):
            if l==0:
                v[0]=(-k*(s**lamb)*u[0]+gammap*u[1])*dt
            elif l==len(u)-1:
                v[len(u)-1]=(k*(s**lamb)*u[len(u)-2])*dt
            elif l==len(u)-2:
                v[len(u)-2]=(-k*(s**lamb)*u[len(u)-2]+k*(s**lamb)*u[len(u)-3]-gammap*(len(u)-2)*u[len(u)-2])*dt
            else:
                v[l]=(-k*(s**lamb)*u[l]+k*(s**lamb)*u[l-1]-gammap*l*u[l]+gammap*(l+1)*u[l+1])*dt
        return v
    tmax=20
    dt=0.0005

    u=np.zeros(int(steps)+1)

    t=0
    count=10
    plim=[]
    sarray=[]
    u[0]=1
    i=0
    s=sb
    while s<100*sb or u[-1]<0.99999:
        u+=du(u=u,sb=sb,t=t)
        t+=dt
        s=sb*np.exp(mu*t)
        k=kk*kd(s=s,H=H,s0h=s0h)
        plim.append(k*(s**lamb)*u[-2]/(mu*s))
        sarray.append(s)
    return np.array(sarray),np.array(plim)

def getk(steps,gammap,s0h,H,lamb,mnsd): 
    """
    Returns the division rate given the parameters and the observed size at division
    
    :Param int Steps: Division steps to trigger division (Positive)
    :Params float gammap: Degradation rate (non-negative)
    :param float s0h: Comitment size (non-negative)
    :param float H: Hill function coefficient (non-negative)
    :param float lamb: exponent of power law for division rate (non-negative)
    :param float mnsd: Mean size at division relative to the mean size at birth (non-negative)
    """ 
    
    
    def get_mean(k):
        """
        Returns the mean added size given the parameters of the function and the division rate k
        :param float k: Division Rate (non-negative)
        """
        sarray,rho= getcum(sb=1,steps=steps,gammap=gammap,s0h=s0h,H=H,lamb=lamb,k=k)
        mnsdt=np.trapz(np.array(rho)*np.array(sarray),x=sarray)
        return mnsdt-mnsd
    k0=1
    m0=get_mean(k0)
    m1=m0
    k1=1
    if m0<0:
        while m0<0:
            k0=k0/2
            m0=get_mean(k0)
            m1=m0
            k1=k0
    while np.sign(m0)==np.sign(m1):
        k1=2*k1
        m1=get_mean(k1)
    #After getting sure that the initial values of k (k0 and k1) predict higher and lower added size
    #it olves the optimal k
    sol = optimize.bisect(get_mean, k0, k1,xtol=1e-5, rtol=1e-5)
    return sol

def getmnsd(sb,steps,gammap,s0h,H,lamb,k):
    """
    Returns the mean added size given the parameters of the function and the division rate k
    :param float k: Division Rate (non-negative)
    """
    sdarr,rhosd=getcum(sb=sb,steps=steps,gammap=gammap,s0h=s0h,H=H,lamb=lamb,k=k)
    mn=np.trapz(np.array(rhosd)*np.array(sdarr),x=np.array(sdarr))
    return mn

def kd(s,H,s0h):
    #Returns the division rate per size unit
    if H==0:
        return 1
    else:
        return 1/(1+(s0h/s)**H)

def getlikebin(data,quantnumber,steps,gammap,s0h,H,lamb,k):
    """
    Estimate the log-likelihood function binning the data using their size at birth.
    :Param pandas DataFrame data: Dataset with columns ['Sb','Sd','Added']
    :Param int quantnumber: Number of quantils for data binning.
    :Param int steps: Division steps to trigger division (Positive).
    :Params float gammap: Degradation rate (non-negative).
    :param float s0h: Comitment size (non-negative).
    :param float H: Hill function coefficient (non-negative).
    :param float lamb: exponent of power law for division rate (non-negative).
    :param float k: Division rate.
    """ 
    like=0
    mnsdd=np.mean(data.Sd)/np.mean(data.Sb)
    
    for i in range(quantnumber):
        lperv0=np.percentile(data.Sb,i*100/quantnumber)
        hperv0=np.percentile(data.Sb,(i+1)*100/quantnumber)
        quanta1=data[data.Sb>lperv0]
        df=quanta1[quanta1.Sb<hperv0]
        sdarr,rhosd=getcum(sb=np.mean(df.Sb)/np.mean(data.Sb),
                           steps=steps,gammap=gammap,s0h=s0h,H=H,lamb=lamb,k=k)

        for sd in df.Sd/np.mean(data.Sb):
            m=0
            while sd>sdarr[m+1]:
                m+=1
            like+=np.log(rhosd[m])
    return like

# %% [code]
def getmnsd(sb,steps,gammap,s0h,H,lamb,k):
    # Estimates the mean size at division given the parameters and the size ath birth sb
    sdarr,rhosd=getcum(sb=sb,steps=steps,gammap=gammap,s0h=s0h,H=H,lamb=lamb,k=k)
    mn=np.trapz(np.array(rhosd)*np.array(sdarr),x=np.array(sdarr))
    return mn

def cv2add(sb,steps,gammap,s0h,H,lamb,k):
    # Estimates the noise in added size given the parameters and the size at birth sb
    sdarr,rhosd=getcum(sb=sb,steps=steps,gammap=gammap,s0h=s0h,H=H,lamb=lamb,k=k)
    ss=np.array(sdarr)
    rhos=np.array(rhosd)
    mn=np.trapz(rhos*ss,x=ss)
    var=np.trapz(rhos*(ss)**2,x=ss)
    CV2=(var-mn**2)/(mn-sb)**2
    return CV2

# %% [code]
def getlikesing(sb,sd,steps,gammap,s0h,H,lamb,k):
    #Estimates the log likelighood of geting a pair (sb,sd) given the parameters
    sdarr,rhosd=getcum(sb=sb,steps=steps,gammap=gammap,s0h=s0h,H=H,lamb=lamb,k=k)
    m=0
    while sd>sdarr[m+1]:
        m+=1
    return np.log(rhosd[m])


def comparemodels(data,stepsopta,ka,stepsoptk,kk,s0h,Hopt,stepsoptg,lambopt,kg,stepsoptr,gammapopt,kr):
    """
    Compare the likelihood of four models given the parameters and the dataset
    :Param pandas DataFrame data: Dataset with columns ['Sb','Sd','Added']
    :Param int stepsopta: Number of division stages for the adder model.
    :Param float ka: Division rate for the adder model.
    :Param int stepsoptk: Number of division stages for the commitment size model.
    :Param float kk: Division rate for the commitment size model.
    :param float s0h: Comitment size (non-negative).
    :param float Hopt: Hill function coefficient (non-negative).
    :Param int stepsoptg: Number of division stages for the commitment size model.
    :Param float lambopt: Exponent for the power law division rate.
    :Param float kg: Division rate for the power law size model.
    :Param float stepsoptr: Division steps for the degradation model.
    :Param float gammapopt: degradation rate.
    :Param float kr: Division rate for the degradation model.
    """     
    pvadd2=data
    sbarrdat=np.array(pvadd2.Sb/np.mean(pvadd2.Sb))
    sdarrdat=np.array(pvadd2.Sd/np.mean(pvadd2.Sb))
    likes=np.empty([len(sbarrdat),3])
    j=0
    for sba in sbarrdat:
        lk=getlikesing(sb=sba,sd=sdarrdat[j],steps=stepsopta,gammap=0,s0h=0,H=0,lamb=1,k=ka)
        likes[j,0]=lk
        lk=getlikesing(sb=sba,sd=sdarrdat[j],steps=stepsoptk,gammap=0,s0h=s0h,H=Hopt,lamb=1,k=kk)
        likes[j,1]=lk
        lk=getlikesing(sb=sba,sd=sdarrdat[j],steps=stepsoptg,gammap=0,s0h=0,H=0,lamb=lambopt,k=kg)
        likes[j,2]=lk
        lk=getlikesing(sb=sba,sd=sdarrdat[j],steps=stepsoptr,gammap=gammapopt,s0h=0,H=0,lamb=1,k=kr)
        likes[j,3]=lk
        j+=1
    likelihood = [np.sum(likes[:,i]) for i in range(len(likes[0]))]
    costo=np.array([1,3,2,2])
    BIC=costo*np.log(len(data))-2*np.array(likelihood)
    AIC=costo*2-2*np.array(likelihood)
    
    pval=np.exp(np.array(likelihood)-np.max(likelihood))
    pval=pval/np.sum(pval)
    
    pvalBIC=np.exp(-(BIC-np.min(BIC)))
    pvalBIC=pvalBIC/np.sum(pvalBIC)
    pvalAIC=np.exp(-(AIC-np.min(AIC)))
    pvalAIC=pvalBIC/np.sum(pvalAIC)
    
    print(f'p-value like:\n Adder={pval[0]}, Commitment={pval[1]}, Power-law={pval[2]}, Degradation={pval[3]}')
    print(f'p-value BIC:\n Adder={pvalBIC[0]}, Commitment={pvalBIC[1]}, Power-law={pvalBIC[2]}, Degradation={pvalBIC[3]}')
    print(f'p-value AIC:\n Adder={pvalAIC[0]}, Commitment={pvalAIC[1]}, Power-law={pvalAIC[2]}, Degradation={pvalAIC[3]}')
    
    print(f'Log-Likelihood: Commitment={likelihood[1]},\n Power-Law={likelihood[2]},\n Degradation={likelihood[3]},\n Adder={likelihood[0]}')
    print(f'BIC: Commitment={round(BIC[1],4)}, Power-Law={round(BIC[2],4)}, Degradation={round(BIC[3],4)}, Adder={round(BIC[0],4)}')
    print(f'AIC: Commitment={round(AIC[1],4)}, Power-Law={round(AIC[2],4)}, Degradation={round(AIC[3],4)}, Adder={round(AIC[0],4)}')

    return np.array(likelihood)

# %% [code]
def filterdata(data,bins):
    #Filters the data outliers
    df=data
    df=df[np.abs(np.log(df.Sb)-np.mean(np.log(df.Sb)))<2.*np.std(np.log(df.Sb))]
    df=df[np.abs(np.log(df.Sb)-np.mean(np.log(df.Sb)))<2.*np.std(np.log(df.Sb))]
    df=df[np.abs(np.log(df.Sd)-np.mean(np.log(df.Sd)))<2.5*np.std(np.log(df.Sd))]
    df=df[np.abs(np.log(df.Sd)-np.mean(np.log(df.Sd)))<2.5*np.std(np.log(df.Sd))]
    pvadd2=df
    df2=pd.DataFrame(columns=pvadd2.columns)
    for i in range(bins):
        lperv0=np.percentile(pvadd2.Sb,i*100/bins)
        hperv0=np.percentile(pvadd2.Sb,(i+1)*100/bins)
        quanta1=pvadd2[pvadd2.Sb>=lperv0]
        df=quanta1[quanta1.Sb<hperv0]
        df=df[np.abs(np.log(df.Added)-np.mean(np.log(df.Added)))<3.5*np.std(np.log(df.Added))]
        df2=pd.concat([df2,df])
    return df2

# %% [code]
"""
Evaluates the models realtive to data.
Input: Model Parameters in './Params.csv' file
Input: Experimental data in './Data.csv' file
Output: Pandas DataFrame dftest with the log-likelihood, AIC, 
BIC and relative likelihood for each model given the data 

"""
params=pd.read_csv('./Params.csv')
data2=pd.read_csv("./Data.csv")
dftest=pd.DataFrame()
for name in data2.Strain.unique():
    df=data2[data2.Strain==name]
    df=filterdata(df,11)
    corr,errcorr=bootcorr(np.array(df.Sb/np.mean(df.Sb)),
                      np.array((df.Sd-df.Sb)/np.mean(df.Sb)))

    print(name,"Corr=(%0.3f +/- %0.3f)"%(corr,errcorr))
    datap=params[params.Strain==name]
    datap=datap.reset_index()
    dic=datap.to_dict()
    mnsd=dic['mnsd'][0]
   
    stepsopta=int(np.mean(df.Added)**2/np.var(df.Added))
    ka=getk(steps=stepsopta,gammap=0,s0h=0,H=0,lamb=1,mnsd=mnsd)
    stepsoptk=dic['M'][2]
    s0h=dic['s0h'][2]
    Hopt=dic['H'][2]
    kk=dic['k'][2]
    stepsoptg=dic['M'][1]
    lambopt=dic['Lamb'][1]
    kg=dic['k'][1]
    stepsoptr=dic['M'][0]
    gammapopt=dic['Gamma'][0]
    kr=dic['k'][0]
    likelihood=comparemodels(data=df,stepsopta=stepsopta,ka=ka,stepsoptk=stepsoptk,kk=kk,s0h=s0h,
              Hopt=Hopt,stepsoptg=stepsoptg,lambopt=lambopt,
              kg=kg,stepsoptr=stepsoptr,gammapopt=gammapopt,kr=kr)
    
    datap['LikeLog']=[likelihood[-1],likelihood[-2],likelihood[-3]]
    costo=np.array([1,3,2,2])
    BIC=costo*np.log(len(df))-2*np.array(likelihood)
    datap['BIC']=[BIC[-1],BIC[-2],BIC[-3]]
    AIC=costo*2-2*np.array(likelihood)
    datap['AIC']=[AIC[-1],AIC[-2],AIC[-3]]    
    pval=np.exp(np.array(likelihood)-np.max(likelihood))
    pval=pval/np.sum(pval)
    pvalBIC=np.exp(-(BIC-np.min(BIC)))
    pvalBIC=pvalBIC/np.sum(pvalBIC)
    datap['pBIC']=[pvalBIC[-1],pvalBIC[-2],pvalBIC[-3]]
    pvalAIC=np.exp(-(AIC-np.min(AIC)))
    pvalAIC=pvalAIC/np.sum(pvalAIC)
    datap['pAIC']=[pvalAIC[-1],pvalAIC[-2],pvalAIC[-3]]  
    datap['Corr']=[corr]*3
    datap['ErrCorr']=[errcorr]*3
    dftest=pd.concat([dftest,datap])

# %% [code]
dftest.to_csv('./Eval.csv',index=False)

# %% [code]
#returns the amount of data of each growth condition
params=pd.read_csv('Params.csv')
data2=pd.read_csv("./Data.csv")

dftest=pd.DataFrame()
for name in data2.Strain.unique():
    df=data2[data2.Strain==name]
    df=filterdata(df,11)
    print(name+':'+str(len(df))+' datapoints')

# %% [code]
#Filters the data especifying a number of bins.
def filterdata(data,bins):
    df=data
    df=df[np.abs(np.log(df.Sb)-np.mean(np.log(df.Sb)))<2.*np.std(np.log(df.Sb))]
    df=df[np.abs(np.log(df.Sb)-np.mean(np.log(df.Sb)))<2.*np.std(np.log(df.Sb))]
    df=df[np.abs(np.log(df.Sd)-np.mean(np.log(df.Sd)))<2.5*np.std(np.log(df.Sd))]
    df=df[np.abs(np.log(df.Sd)-np.mean(np.log(df.Sd)))<2.5*np.std(np.log(df.Sd))]
    pvadd2=df
    df2=pd.DataFrame(columns=pvadd2.columns)
    for i in range(bins):
        lperv0=np.percentile(pvadd2.Sb,i*100/bins)
        hperv0=np.percentile(pvadd2.Sb,(i+1)*100/bins)
        quanta1=pvadd2[pvadd2.Sb>=lperv0]
        df=quanta1[quanta1.Sb<hperv0]
        df=df[np.abs(np.log(df.Added)-np.mean(np.log(df.Added)))<3.5*np.std(np.log(df.Added))]
        df2=pd.concat([df2,df])
    return df2

# %% [code]
#Plots the data moments and the predictions from the models

from matplotlib.offsetbox import AnchoredText
fig, ax = plt.subplots(2,8, figsize=(12,2.5),sharex=True)
data2=pd.read_csv("./Data.csv")
namearr=['Str1','Str2','Str3','Str4','Str5','Str6','Str7','Str8']
table=[]

dataN=pd.read_csv('./NSol.csv')
for j in range(len(namearr)):
    name=namearr[j]
    pvadd2=data2[data2.Strain==name]
    pvadd2=filterdata(pvadd2,11)
    slope, intercept, r_value, p_value, std_err = stats.linregress(pvadd2.Sb/np.mean(pvadd2.Sb),
                                                            (pvadd2.Sd-pvadd2.Sb)/np.mean(pvadd2.Sb))
    
    corr,errcorr=bootcorr(np.array(pvadd2.Sb/np.mean(pvadd2.Sb)),
                          np.array((pvadd2.Sd-pvadd2.Sb)/np.mean(pvadd2.Sb)))

    print(name,"Corr=(%0.3f +/- %0.3f)"%(corr,errcorr))
    sbarr=[]
    deltarr=[]
    allsb=pvadd2.Sb.tolist()
    alldelt=(pvadd2.Sd-pvadd2.Sb).tolist()
    for i in range(1000):
        r=np.random.randint(len(allsb))
        sbarr.append(allsb[r])
        deltarr.append(alldelt[r])
    ax[0][j].scatter(np.array(sbarr)/np.mean(pvadd2.Sb),np.array(deltarr)/np.mean(pvadd2.Sb),s=0.5,c='#319AD8')
    

    CV2d=[]
    delt=[]
    sbarr=[]
    
    errcv2d=[]
    errdelt=[]
    errsb=[]
    quantnumber=5
    bins=np.linspace(np.min(pvadd2.Sb),np.max(pvadd2.Sb),quantnumber+1)
    for i in range(quantnumber):
        quanta1=pvadd2[pvadd2.Sb>bins[i]]
        quanta2=quanta1[quanta1.Sb<bins[i+1]]
        
        mean_cntr, var_cntr, std_cntr = bayesest((quanta2.Added)/np.mean(pvadd2.Sb),alpha=0.95)
        meanv0_cntr, varv0_cntr, stdv0_cntr = bayesest((quanta2.Sb)/np.mean(pvadd2.Sb),alpha=0.95)
        CV2d.append(np.var((quanta2.Added)/np.mean(pvadd2.Sb))/
                    np.mean((quanta2.Added)/np.mean(pvadd2.Sb))**2)
        delt.append(np.mean((quanta2.Added)/np.mean(pvadd2.Sb)))
        sbarr.append(np.mean(quanta2.Sb)/np.mean(pvadd2.Sb))
        
        errv=((var_cntr[1][1]-var_cntr[0])/mean_cntr[0]**2)**2+(2*(mean_cntr[1][1]-mean_cntr[0])*var_cntr[0]/mean_cntr[0]**3)**2
    
        errcv2d.append(np.sqrt(errv))
        errdelt.append(mean_cntr[1][1]-mean_cntr[0])
        errsb.append(meanv0_cntr[1][1]-meanv0_cntr[0])

    ax[0][j].errorbar(np.array(sbarr),np.array(delt),xerr=errsb,yerr=errdelt, fmt='o',mec='#094264',
                      capsize=3,markersize='3',elinewidth=1,c='#0075BD')
    ax[1][j].errorbar(np.array(sbarr),100*np.array(CV2d),xerr=errsb,yerr=100*np.array(errcv2d),
                      fmt='o',mec='#094264',
                      capsize=3,markersize='3',elinewidth=1,c='#0075BD')
    
    slope, intercept, r_value, p_value, std_err = stats.linregress(pvadd2.Sb/np.mean(pvadd2.Sb),(pvadd2.Sd-pvadd2.Sb)/np.mean(pvadd2.Sb))
    xarr=np.linspace(np.min(pvadd2.Sb/np.mean(pvadd2.Sb)),np.max(pvadd2.Sb/np.mean(pvadd2.Sb)),100)
    var=np.var((pvadd2.Sd-pvadd2.Sb)/np.mean(pvadd2.Sb))
    ax[1][j].set_ylim(100*0.7*np.min(CV2d),100*1.3*np.max(CV2d))   
    ax[1][j].set_xlim(0.6,1.5)
    
    nstrain=dataN[dataN.Strain==name]
    ndeg=nstrain[nstrain.Model=='Deg']
    ax[0][j].plot(ndeg.Sb,ndeg.Added,'r',ls=':',lw=1.5)
    ax[1][j].plot(ndeg.Sb,100*ndeg.CV2D,'r',ls=':',lw=1.5)
    
    ndeg=nstrain[nstrain.Model=='Pow']
    ax[0][j].plot(ndeg.Sb,ndeg.Added,'g',ls='--',lw=1.5)
    ax[1][j].plot(ndeg.Sb,100*ndeg.CV2D,'g',ls='--',lw=1.5)  
    
    ndeg=nstrain[nstrain.Model=='Comm']
    ax[0][j].plot(ndeg.Sb,ndeg.Added,'k',lw=1.5)
    ax[1][j].plot(ndeg.Sb,100*ndeg.CV2D,'k',lw=1.5) 
    taqui=100*np.arange(np.round(0.8*np.min(CV2d),2),1.2*np.max(CV2d),step=0.01)
    ax[1][j].set_yticks(taqui) 
    ax[0][j].set_yticks([0.5,1.0,1.5]) 
    if j==7:
        ax[0][j].set_yticks([2,3,4]) 
        ax[0][j].set_ylim(1.5,4.5)
    setaxis(ax[0,j])
    setaxis(ax[1,j])
    plt.subplots_adjust(wspace=0.35)
    ax[1,j].set_xlabel(r'$s_b/\langle s_b \rangle$')
    at = AnchoredText(
    "N=%d"%(len(pvadd2)), prop=dict(size=7), frameon=True, loc='upper right')
    at.patch.set_boxstyle("round,pad=0.,rounding_size=0.2")
    ax[0,j].add_artist(at)
    ax[0,j].set_title('Str'+str(j+1),fontsize=10)
ax[0,0].set_ylabel(r'$\Delta/\langle s_b \rangle$')
ax[1,0].set_ylabel(r'$CV^2_\Delta$')
ax[1,0].text(0.3,3.2,r'$\times 1/0^{-2}$',fontsize=8)
#plt.savefig('./noisesig.eps',bbox_inches='tight',dpi=400)

# %% [code]
#Return the numerical solution given the parameters in './Params.csv'
fig, ax = plt.subplots(2,8, figsize=(24,5),sharex=True)
data2=pd.read_csv("./Data.csv")

namearr=['Str1','Str2','Str3','Str4','Str5','Str6','Str7','Str8']
table=[]

params=pd.read_csv('./Params.csv')
numberdata=pd.DataFrame()
for j in tqdm(range(len(namearr))):
    name=namearr[j]
    pvadd2=data2[data2.Medium==name]
    pvadd2=filterdata(pvadd2,11)
    sbarr=[]
    deltarr=[]
    allsb=pvadd2.Sb.tolist()
    alldelt=(pvadd2.Sd-pvadd2.Sb).tolist()
    for i in range(1000):
        r=np.random.randint(len(allsb))
        sbarr.append(allsb[r])
        deltarr.append(alldelt[r])
    ax[0][j].scatter(np.array(sbarr)/np.mean(pvadd2.Sb),np.array(deltarr)/np.mean(pvadd2.Sb),s=1.5)
    quantnumber=7
    bins=np.linspace(np.min(pvadd2.Sb),np.max(pvadd2.Sb),quantnumber)

    CV2d=[]
    delt=[]
    sbarr=[]
    
    errcv2d=[]
    errdelt=[]
    errsb=[]
    for i in range(quantnumber):
        quanta1=pvadd2[pvadd2.Sb>bins[np.max([0,i-1])]]
        quanta2=quanta1[quanta1.Sb<bins[np.min([len(bins)-1,i+1])]]

        
        mean_cntr, var_cntr, std_cntr = bayesest((quanta2.Added)/np.mean(pvadd2.Sb),alpha=0.95)
        
        meanv0_cntr, varv0_cntr, stdv0_cntr = bayesest((quanta2.Sb)/np.mean(pvadd2.Sb),alpha=0.95)
        
        CV2d.append(np.var((quanta2.Added)/np.mean(pvadd2.Sb))/
                    np.mean((quanta2.Added)/np.mean(pvadd2.Sb))**2)
        delt.append(np.mean((quanta2.Added)/np.mean(pvadd2.Sb)))
        sbarr.append(np.mean(quanta2.Sb)/np.mean(pvadd2.Sb))
        
        errv=((var_cntr[1][1]-var_cntr[0])/mean_cntr[0]**2)**2+(2*(mean_cntr[1][1]-mean_cntr[0])*var_cntr[0]/mean_cntr[0]**3)**2
    
        errcv2d.append(np.sqrt(errv))
        errdelt.append(mean_cntr[1][1]-mean_cntr[0])
        errsb.append(meanv0_cntr[1][1]-meanv0_cntr[0])


    ax[0][j].errorbar(np.array(sbarr),np.array(delt),xerr=errsb,yerr=errdelt, fmt='o',mec='k',
                      capsize=5,markersize='5',elinewidth=1.5,c='#0075BD')
    ax[1][j].errorbar(np.array(sbarr),CV2d,xerr=errsb,yerr=errcv2d, fmt='o',mec='k',
                      capsize=5,markersize='5',elinewidth=1.5,c='#0075BD')
    
    datap=params[params.Strain==name]
    datap=datap.reset_index()
    dic=datap.to_dict()
    mnsd=dic['mnsd'][0]
   
    stepsopta=int(np.mean(pvadd2.Added)**2/np.var(pvadd2.Added))
    ka=getk(steps=stepsopta,gammap=0,s0h=0,H=0,lamb=1,mnsd=mnsd)
    stepsoptk=dic['M'][2]
    s0hopt=dic['s0h'][2]
    Hopt=dic['H'][2]
    kk=dic['k'][2]
    sbdet=np.linspace(0.3,1.8,15)
    sddet=[getmnsd(sb=sb,steps=stepsoptk,gammap=0,s0h=s0hopt,H=Hopt,lamb=1,k=kk) for sb in sbdet]
    CV2det=[cv2add(sb=sb,steps=stepsoptk,gammap=0,s0h=s0hopt,H=Hopt,lamb=1,k=kk) for sb in sbdet]
    dic2={'Strain':[name]*len(sbdet),'Model':['Comm']*len(sbdet),'Gamma':[0]*len(sddet),'Lamb':[1]*len(sddet),
         's0h':[s0hopt]*len(sddet),'H':[Hopt]*len(sddet),'M':[stepsoptk]*len(sddet),'k':[kk]*len(sddet),
         'Sb':sbdet,'Added':np.array(sddet)-np.array(sbdet),'CV2D':CV2det}

    df=pd.DataFrame(dic2)
    numberdata=pd.concat([numberdata,df])


    ax[0][j].plot(sbdet,sddet-sbdet,'k',lw=1)
    ax[1][j].plot(sbdet,CV2det,'k',lw=1)
    
    stepsoptg=dic['M'][1]
    lambopt=dic['Lamb'][1]
    kg=dic['k'][1]
    sddet=[getmnsd(sb=sb,steps=stepsoptg,gammap=0,s0h=0,H=0,lamb=lambopt,k=kg) for sb in sbdet]
    CV2det=[cv2add(sb=sb,steps=stepsoptg,gammap=0,s0h=0,H=0,lamb=lambopt,k=kg) for sb in sbdet]


    ax[0][j].plot(sbdet,sddet-sbdet,'g',lw=1)
    ax[1][j].plot(sbdet,CV2det,'g',lw=1)
    dic2={'Strain':[name]*len(sbdet),'Model':['Pow']*len(sbdet),'Gamma':[0]*len(sddet),'Lamb':[lambopt]*len(sddet),
         's0h':[0]*len(sddet),'H':[0]*len(sddet),'M':[stepsoptg]*len(sddet),'k':[kg]*len(sddet),
         'Sb':sbdet,'Added':np.array(sddet)-np.array(sbdet),'CV2D':CV2det}
    df=pd.DataFrame(dic2)
    numberdata=pd.concat([numberdata,df])
    stepsoptr=dic['M'][0]
    gammapopt=dic['Gamma'][0]
    kr=dic['k'][0]
    sddet=[getmnsd(sb=sb,steps=stepsoptr,gammap=gammapopt,s0h=0,H=0,lamb=1,k=kr) for sb in sbdet]
    CV2det=[cv2add(sb=sb,steps=stepsoptr,gammap=gammapopt,s0h=0,H=0,lamb=1,k=kr) for sb in sbdet]


    ax[0][j].plot(sbdet,sddet-sbdet,'r',lw=1)
    ax[1][j].plot(sbdet,CV2det,'r',lw=1)    
    
    dic2={'Strain':[name]*len(sbdet),'Model':['Deg']*len(sbdet),'Gamma':[gammapopt]*len(sddet),'Lamb':[1]*len(sddet),
         's0h':[0]*len(sddet),'H':[0]*len(sddet),'M':[stepsoptr]*len(sddet),'k':[kr]*len(sddet),
         'Sb':sbdet,'Added':np.array(sddet)-np.array(sbdet),'CV2D':CV2det}

    df=pd.DataFrame(dic2)
    numberdata=pd.concat([numberdata,df])
    
    
    slope, intercept, r_value, p_value, std_err = stats.linregress(pvadd2.Sb/np.mean(pvadd2.Sb),(pvadd2.Sd-pvadd2.Sb)/np.mean(pvadd2.Sb))
    xarr=np.linspace(np.min(pvadd2.Sb/np.mean(pvadd2.Sb)),np.max(pvadd2.Sb/np.mean(pvadd2.Sb)),100)
    var=np.var((pvadd2.Sd-pvadd2.Sb)/np.mean(pvadd2.Sb))
    ax[1][j].set_ylim(0.7*np.min(CV2d),1.3*np.max(CV2d))   
    ax[1][j].set_xlim(0.6,1.5)
    taqui=np.arange(np.round(0.7*np.min(CV2d),2),1.3*np.max(CV2d),step=0.01)
    ax[1][j].set_yticks(taqui) 
    
 
    setaxis(ax[0,j])
    setaxis(ax[1,j])
    plt.subplots_adjust(wspace=0.3)

# %% [code]
numberdata.to_csv('./NSol.csv',index=False)
```
