## Supplementary material for "Mechanisms of Cell Size Regulation in Slow-Growing *Escherichia coli* Cells: Discriminating Models Beyond the Adder": README.md

# commitment-size
@author: canietoa. Cesar Augusto Nieto Acuna. University of Delaware 2023.

Source code for the article:  Mechanisms of Cell Size Regulation in Slow-Growing Escherichia coli Cells: Discriminating Models Beyond the Adder

Datasets:

Data.csv: Contains the Size at both and size at division for different strains following the sizer-like division strategy.

Eval2.csv: Evaluation of the models performance given the parameters and the data. 

NSol.csv: Results of the numerical estimation of the moments of the distribution of size at division gven the size at birth.

Params.csv: Best model parameters for the observed data.

Code:

DataAnalysis.ipynb: Code for the estimation of distributions and model evaluation. Also presents how the main figure was plotted. 
